# Tracing the dynamic recruitment trajectory of Optineurin between the centrosome and other macromolecular assemblies, signifying its role in cell cycle progression

**DOI:** 10.64898/2026.09.30.755612

**Authors:** Ankan Chakraborty, Koyena Kundu, Dominique Weil, Bishwajit Kundu

## Abstract

Spatiotemporal allocation and dynamic redistribution of centrosomal proteins are critical for coordinating the structural and molecular events that ensure successful cell division. While depletion of the multifunctional protein Optineurin (OPTN) impairs cell-cycle progression, our previous report of the presence of OPTN at key cell-division assemblies raised the question of how its trafficking is coordinated during cell division. In this study, using cell-cycle synchronisation coupled with flow cytometric analysis and confocal microscopy, we report that OPTN progressively dissociates from centrosomes during mitosis, redistributes to cytokinetic structures, accumulates at the spindle midzone, and then concentrates at the midbody during cytokinesis. Moreover, following this transition, OPTN progressively re-emerges at centrosomes during G1 of the subsequent cell cycle. Further, microtubule perturbation and regrowth assays provide evidence that OPTN recruitment and retention at the pericentriolar matrix (PCM) can occur independently of microtubules, whereas its exit is microtubule-dependent. Collectively, this study depicts, for the first time, a detailed movement trail of OPTN, highlighting its centrosome-linked redistribution and microtubule dependency. This context-dependent localisation and trafficking of OPTN across the centrosome-midbody axis establish a definitive spatiotemporal roadmap that can be strategically utilised to study its functional role in cell division.

Illustration of OPTN trafficking across the centrosome-midbody axis during cell cycle progression (A) and the regulated OPTN’s centrosomal entry and exit (B).

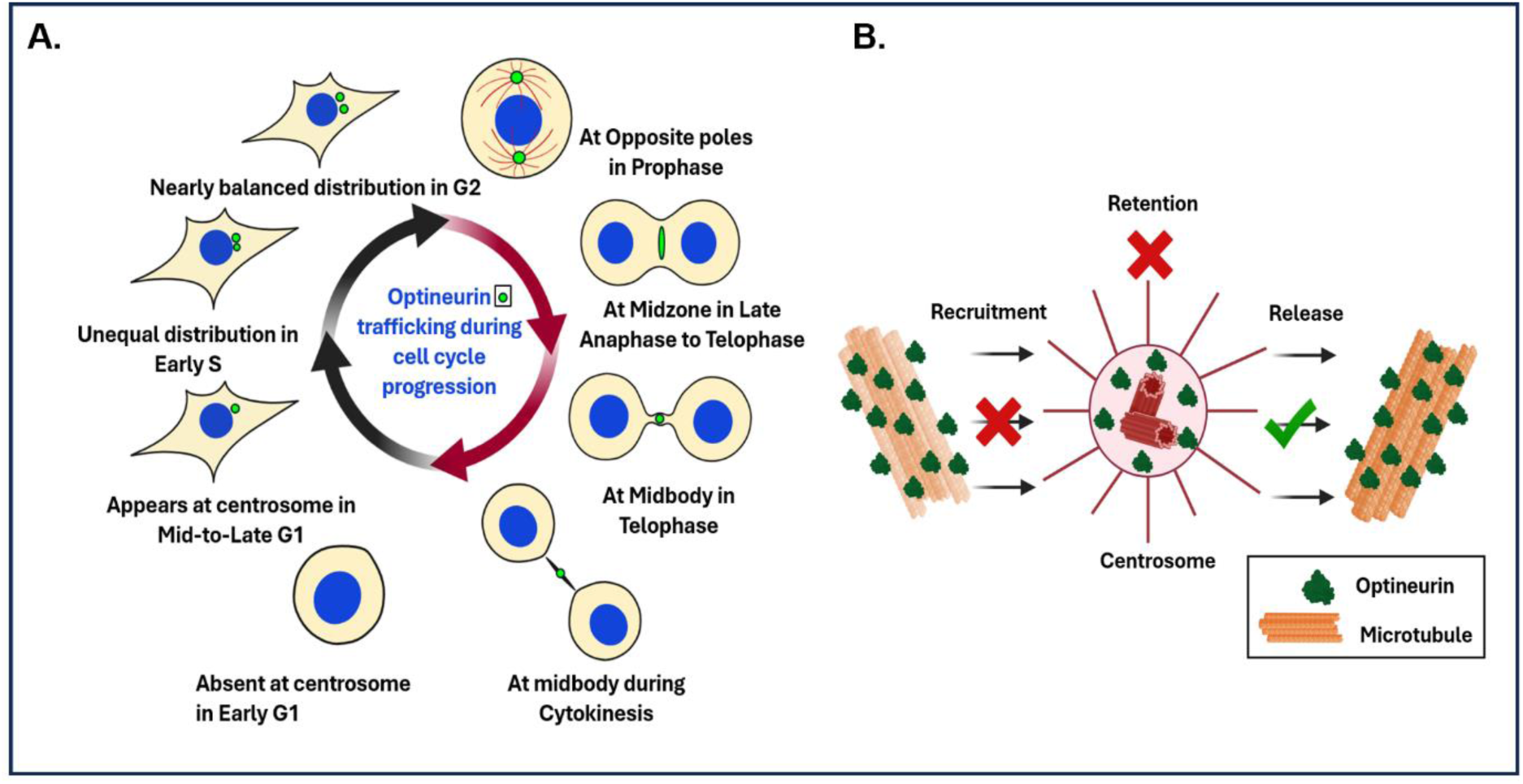

## Introduction

In animal cells, centrosomes act as the primary microtubule-organising centres (MTOCs) ^1,2^. Centrosomes participate in an active exchange of components with the cytoplasm, enabling the dynamic yet precisely regulated assembly of the spindle and segregation of chromosomes ^3–5^. The coordinated protein redistribution process continues past the chromosome segregation event, where several proteins, including the centrosomal protein of 55 kDa (CEP55) and Centriolin, gradually leave the centrosome and temporarily relocate to the midbody during cytokinesis ^6,7^. Disrupting this coordinated trafficking can severely impair cytokinesis, compromise cell division, and increase the risk of aneuploidy.

Orchestrated protein trafficking continues into the subsequent cell cycle, re-establishing the designated centrosomal architecture with a mature PCM ^8,9^. Following cytokinesis, as the newly formed daughter cells re-enter early G1 and progress through the S phase, the centriole pair undergoes a tightly controlled cycle of disengagement and maturation, respectively ^10^. Synchronised with centriolar proximity and re-attachment, the PCM also reassembles and matures through iterative recruitment, retention and release/exit of numerous proteins, restoring a fully functional, remodelled centrosome ^9^. This remodelling involves protein recruitment at the centrosome through a) microtubule-dependent transport and b) local scaffold-mediated assembly ^11–14^. Active exit routes ensure balanced redistribution of proteins to other cellular locations when needed ^15^. Hindering this balance can significantly affect centrosomal structure, compromising cell division.

OPTN is a microtubule-associated protein involved in diverse cellular processes ^16–18^. Its extensive interaction network links it to vesicular trafficking, innate immune signalling, and various forms of autophagy, particularly aggrephagy and mitophagy ^16,19–22^. Emerging evidence also suggests a role for OPTN in cell cycle control, in which Golgi-associated OPTN interacts with polo-like kinase 1 (Plk1) and translocates to the nucleus, where it inactivates Plk1 and regulates proper mitotic progression ^23^. Importantly, mutants of OPTN are linked with several neurodegenerative disorders (NDs), including amyotrophic lateral sclerosis (ALS) and primary open-angle glaucoma (POAG); however, the connection between OPTN’s cell-cycle-associated functions and neurodegeneration remains poorly understood ^24,25^. As defective mitosis and aberrant cell division have been increasingly linked to NDs, understanding how OPTN participates in cell cycle progression becomes especially relevant ^26^.

OPTN’s ubiquitous presence, together with our previous findings of its localisation at the centrosome, spindle, and midbody during the cell cycle, catalysed a focused research initiative to understand OPTN’s dynamic trafficking ^27^. However, since our previous study was largely based on an asynchronous cell population, we had limited resolution of OPTN’s temporal dynamics across cell-cycle stages. In this study, we address these research gaps by combining synchronised cell populations, confocal microscopy, and targeted perturbation of the microtubule cytoskeleton. We define a foundational framework for a cell cycle-dependent centrosome-linked trafficking pathway for OPTN, encompassing its release from centrosomes during mitosis, redistribution to cytokinetic structures, and re-establishment at centrosomes in the subsequent cell cycle. In addition, we provide the first detailed analysis of the spatial organisation of OPTN within the centrosome in relation to the microtubule network. Given the complexity and diversity of cell division assemblies, identifying the precise sites of OPTN’s functionality remains challenging. Here, our study revealed the dynamic trafficking of OPTN and its regulated centrosomal recruitment and exit, providing a comprehensive framework for dissecting its context-dependent function during cell cycle progression.

## Results

### 1. OPTN progressively dissociates from the centrosome during mitotic progression

To investigate the dynamic connection of OPTN with the centrosome during mitosis, HeLa cells were subjected to a double thymidine block and released into new media to synchronise the cells into mitosis (Fig. 1a). We chose HeLa cells for the following reasons that they are a well-established mammalian cell model for cell-cycle studies, they possess unambiguous centrosomal architecture, and they can be efficiently synchronised and quantitatively analysed for centrosomal characteristics. Thymidine is a DNA synthesis inhibitor that arrests cells at the G1/S boundary prior to DNA replication ^28^. A double thymidine block was therefore used to efficiently synchronise cells at the G1/S boundary before release for subsequent cell-cycle progression. For quantitative analysis, synchronised cells were tracked for a span of 2 hours beginning from 10 h to 12 h post-release, encompassing the full progression from prophase through cytokinesis (Fig. 1a). Cell cycle stages were confirmed by flow cytometry analysis (Fig. 1b) at each time point, from release, to track the mitotic stages as shown in Fig. 1c. Cells were co-stained for OPTN and γ-tubulin (a centrosomal marker) and categorised into five groups based on mitotic stage: prophase/prometaphase, metaphase, anaphase I, anaphase II, telophase/cytokinesis ^29^. Immunofluorescence analysis revealed that OPTN was associated with centrosomes during early mitotic stages. However, measuring the OPTN:γ-tubulin corrected mean intensity ratio at the γ-tubulin-positive centrosomes revealed a gradual reduction of OPTN from anaphase I to anaphase II, with barely detectable signal by the end of telophase/cytokinesis, when OPTN was observed at the midbody during cytokinesis (Fig. 1d, 1e).

**Fig. 1.**
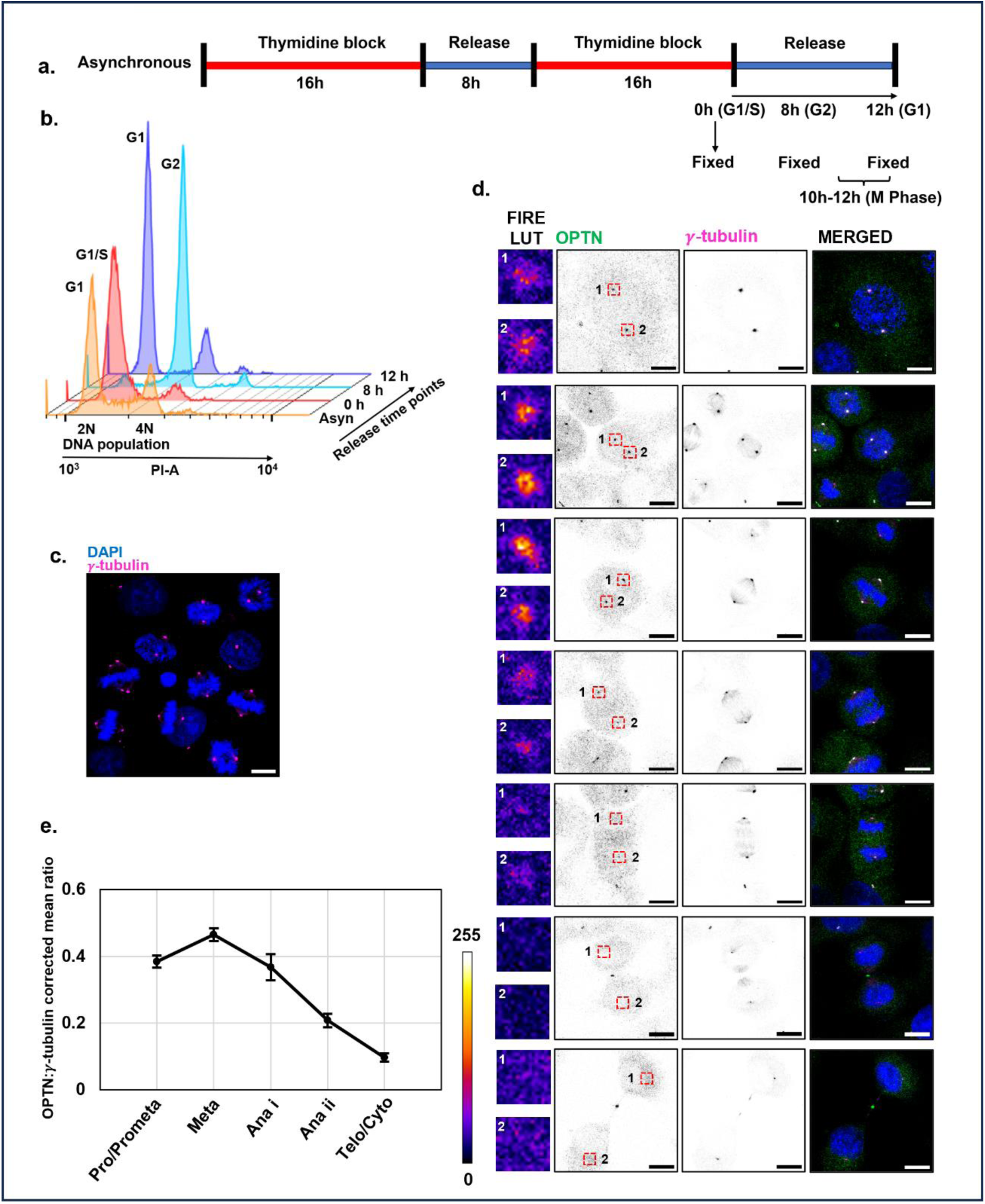
Cell cycle-dependent reduction of OPTN signal from the centrosome during mitosis in HeLa cells. (a) Schematic representation of a double-thymidine block following release to get an enriched population of cells in the mitotic phase. (b) Flow cytometry profiles of propidium iodide (PI)-stained HeLa cells following double thymidine block and subsequent time points during release. (c) An enriched population of metaphase cells after synchronisation. (d) Representative confocal images of cells from each mitotic population showing OPTN (green), γ-tubulin (magenta), and the merged channel with 4′,6-diamidino-2-phenylindole (DAPI)-stained (blue) nucleus. Single channels are shown as inverted grayscale images. OPTN shows strong centrosomal localisation during prophase/metaphase, which progressively diminishes during anaphase II and becomes nearly undetectable during cytokinesis. Fire look-up table (LUT) (leftmost column) representation of centrosomal OPTN intensity (marked as 1 and 2), highlighting a gradual reduction of centrosomal OPTN as cells progress through mitosis. (e) OPTN signal at the centrosome was quantified for individual centrosomes as the ratio of corrected mean OPTN fluorescence to corrected mean γ-tubulin fluorescence (OPTN_corrected mean_: γ-tubulin_corrected mean_). The data represent the mean ± standard error of the mean (SEM) across n=18-42 centrosomes for each stage. Scale bars, 10 µm.

To further explore trafficking patterns, we examined OPTN dynamics in noncancerous HEK293 cells. We performed a double thymidine block and subsequent cell release to obtain an enriched mitotic population. We tracked the synchronised cells for 1 h, from 11 h to 12 h post-release (Fig. 2a, 2b). We observed that OPTN gradually decreased at the centrosome as the cell cycle progressed through mitosis. While it became nearly undetectable at the centrosome, we observed a prominent OPTN signal localised at the midbody during cytokinesis (Fig. 2c, 2d). This pattern resembles observations in HeLa cells, indicating similar OPTN trafficking across cell types.

**Fig. 2.**
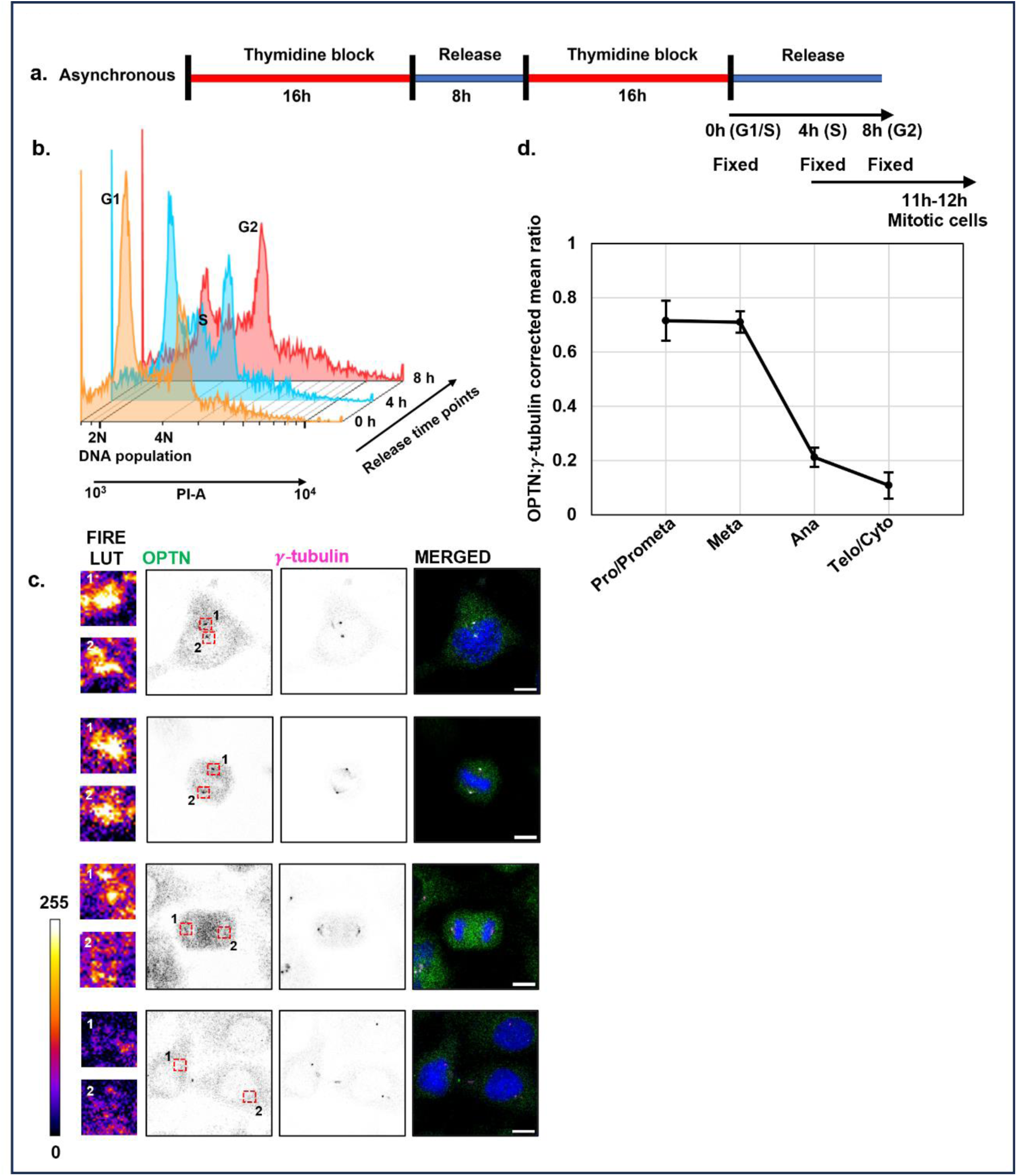
Cell cycle-dependent reduction of OPTN signal from the centrosome during mitosis in HEK 293 cells. (a) Schematic representation of a double-thymidine block following release to get an enriched population of cells in the mitotic phase. (b) Flow cytometry profiles of propidium iodide (PI)-stained HEK 293 cells following a double thymidine block and subsequent time points during release. (c) Representative confocal images of cells from each mitotic population showing OPTN (green), γ-tubulin (magenta), and the merged channel with a DAPI-stained (blue) nucleus. Single channels are shown as inverted grayscale images. OPTN shows strong centrosomal localisation during prophase/metaphase, which progressively diminishes during anaphase and becomes nearly undetectable during cytokinesis. Fire look-up table (LUT) (leftmost column) representation of centrosomal OPTN intensity (marked as 1 and 2), highlighting a gradual reduction of centrosomal OPTN as cells progress through mitosis. (d) OPTN signal at the centrosome was quantified for individual centrosomes as the ratio of corrected mean OPTN fluorescence to corrected mean γ-tubulin fluorescence (OPTN_corrected mean_: γ-tubulin_corrected mean_). The data represent the mean ± SEM across n=18-40 centrosomes for each stage. Scale bars, 10 µm.

### 2. OPTN redistributes to the spindle midzone and contractile ring during cytokinesis

During cytokinesis, OPTN localisation at the midbody, with minimal signal at the centrosome in both HeLa and HEK 293 cell lines, suggests a possible relocation of the OPTN pool from the centrosome to the midbody. We investigated whether this relocalisation follows intermediate, transient associations with other mitotic structures before reaching the midbody. We examined synchronised HeLa cells by the thymidine-mediated blocking and release experiment as described above (see results, section 1) (Fig. 3a). We observed an accumulation of OPTN at the spindle midzone during late anaphase to telophase (Fig. 3b). To understand its spatial relationship with the cytokinetic machinery, cells were co-stained with non-muscle myosin IIB (NMIIB), a marker of the contractile ring ^30^. Confocal microscopy and quantitative analysis revealed that early in telophase, OPTN formed an inner disc-like structure at the midzone, surrounded by the ring-shaped NMIIB (Fig. 3c, 3d, 3e, 3f). As cytokinesis progressed, OPTN became more confined and eventually localised at the midbody junction (Fig. 3b). These findings demonstrate that, after detaching from the centrosome, OPTN is redistributed to the midzone, gradually becomes confined within the contractile ring, and finally localises at the midbody during cytokinesis.

**Fig. 3.**
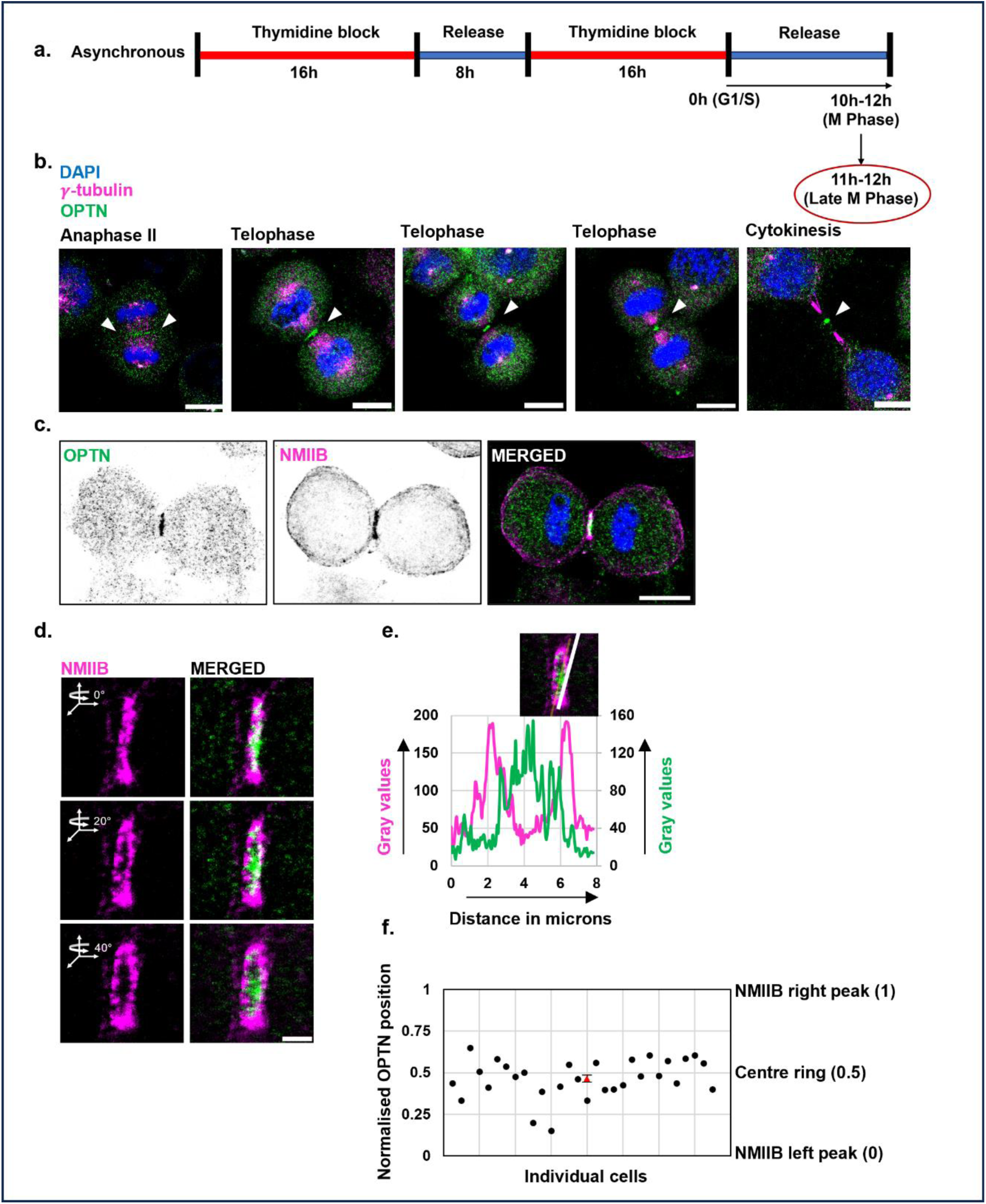
Recruitment of OPTN to the spindle midzone and midbody during cytokinesis. (a) Schematic representation of the double-thymidine block following release to get an enriched population of HeLa cells with late-mitotic (M) phases. (b) Representative confocal images of cells from each mitotic population, co-stained for OPTN (green) and γ-tubulin (magenta), showing a gradual transition of OPTN from the spindle midzone to the midbody as cells progress from telophase to cytokinesis. The merged image shows three channels: OPTN (green), NMIIB (magenta), and DAPI (blue). Arrowheads indicate OPTN at the midzone and midbody. The images shown at the indicated cell-cycle stages were obtained from fixed cells and represent different cells at each stage, rather than real-time imaging of the same cell progressing through mitosis. (c) Co-immunostaining of OPTN (green) and NMIIB (magenta) reveals a solid, core-like OPTN structure at the midzone surrounded by the ring-shaped NMIIB contractile ring. The merged image shows three channels: OPTN (green), NMIIB (magenta), and DAPI (blue). Single channels are shown as inverted grayscale images. (d) Three-dimensional reconstruction of confocal z-stacks illustrating spatial confinement of OPTN (green) to the central core of the midzone/midbody (magenta) region. The merged image shows two channels: OPTN (green) and NMIIB (magenta). For (b) and (c), the scale bars are 10 µm and for (d) the scale bar is 2µm. (e) Line-scan intensity profiles across the cytokinetic plane (at 40° rotation) show a central OPTN peak (green) flanked by peripheral NMIIB (magenta) signals, confirming core-versus-ring organisation. (f) For each line scan, the two NMIIB intensity peaks defining the boundaries of the contractile ring were normalised to a 0-1 scale, with 0 and 1 representing the left and right peaks, respectively. The position of the OPTN intensity peak was plotted on this normalised scale to determine its spatial distribution within the NMIIB ring, with 0.5 representing the centre of the ring. Individual cells are shown as scatter points, with the mean ± SEM indicated by a red triangle (n=30 cells).

### 3. OPTN reappears at the centrosome during G1 of the subsequent cell cycle

Given that OPTN becomes undetectable at centrosomes during cytokinesis, we next investigated when centrosomal OPTN is re-established following mitotic exit. To address this, HeLa cells were synchronised at the G2/M boundary using nocodazole, a microtubule-depolymerising agent that disrupts spindle microtubules and arrests cells at the G2/M transition ^31,32^. The arrested cells were subsequently collected by shake-off, replated in fresh medium, and released from nocodazole treatment to allow synchronous progression through mitosis and entry into the subsequent cell cycle (Fig. 4a). Following release, cells were collected at defined time points and analysed by flow cytometry to determine cell cycle stages (Fig. 4b). Analysis revealed that by 3 h post-release, most cells exited mitosis and entered the G1 phase of the next cell cycle (Fig. 4b). Immunofluorescence analysis was performed at this time point for OPTN and γ-tubulin. In agreement with previous reports, a substantial number of centrosomes with split centrioles were detected at 3 h post-release (early G1) ^10^. For our analysis, split centrosomes were defined as those at the early G1 stage, with a distance between the two centrioles greater than 0.5 µm. As the cell cycle progressed, these gradually came into proximity (inter-centriolar distance ≤0.5 µm) and/or underwent apparent fusion into a single centrosome (Fig. 4c, 4d). Notably, we observed OPTN-positive centrosomes predominantly at 12 h post-release (Fig. 4c, 4e). Together, these results suggest that OPTN does not immediately localise to the immature split centrioles in early G1 but gets re-recruited to the centrosome as the cell cycle progresses.

**Fig. 4.**
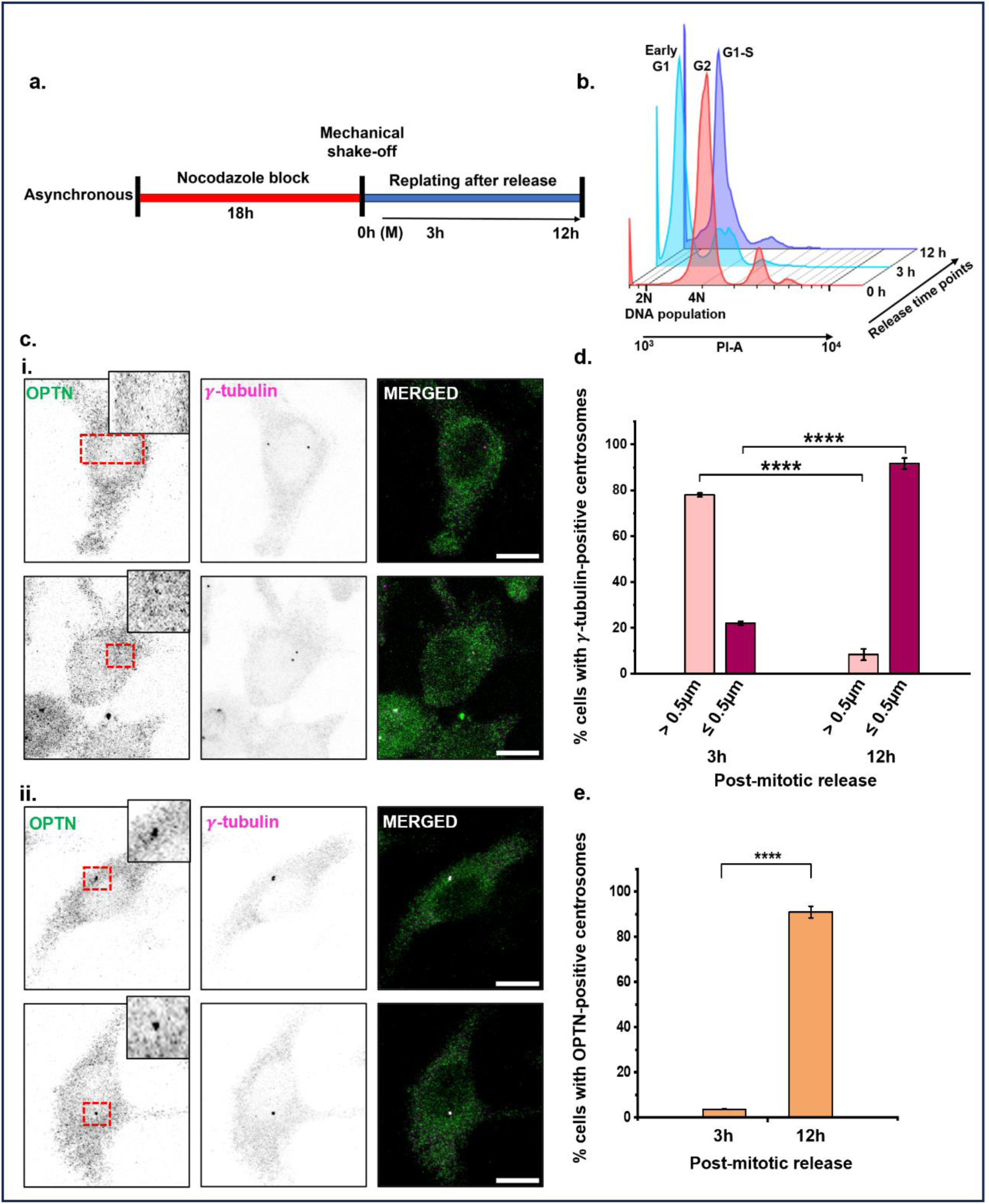
Re-establishment of centrosomal OPTN during G1 of the next cell cycle. (a) Schematic representation of the nocodazole block following release to get an enriched population of cells in mitotic phases (M). (b) Flow cytometry profiles of propidium iodide (PI)-stained HeLa cells following nocodazole block and release at subsequent time points. (c) Confocal images of cells after release into G1, stained for OPTN (green) and γ-tubulin (magenta), were classified into two groups: (i) cells with split centrosomes (Early G1, inter-centriolar distance >0.5 µm), and (ii) cells with centrosomes where centrioles were in proximity (inter-centriolar distance ≤0.5 µm) or appeared as a single centrosome. Single channels are shown as inverted grayscale images. Zoomed views of the centrosomal area (red inset) are shown in the upper right corner of each image of the green channel. (d) The percentage of cells at different time points (3h and 12h) as cells entered G1 was quantified for these two classified groups (c. i. and c. ii.) and plotted graphically, showing a gradual increase in group ii centrosomes as cells progressed through G1. (e) Quantitative estimate showing an increasing number of cells with OPTN-positive centrosome with time. Data represent mean ± SEM for three independent events with 20-33 centrosomes analysed per experiment. Statistical analysis was performed using an unpaired two-tailed *t-test*, with significance defined as *p < 0.05, **p < 0.01, ***p < 0.001, ****p < 0.0001. Scale bars, 10 µm.

### 4. OPTN shows stage-specific preferential distribution between mother and daughter centrioles at the G1/S boundary

We previously reported a preferential distribution of OPTN between the two centrioles in asynchronous cell populations. To precisely define the timing and dynamics of this unequal distribution, we performed a quantitative analysis using synchronised cells. HeLa cells were synchronised at the G1/S boundary using a double thymidine block and released into fresh medium to catch the subsequent S phase, which was tracked by flow cytometry (Fig. 5a, 5b). OPTN asymmetry between two centrioles was calculated as a percentage using the formula 100 × [(H − L)/H], where H and L represent the higher and lower OPTN total intensities [corrected total cell fluorescence (CTCF)], respectively, measured at the two centrioles. The percentage of cells exhibiting an unequal distribution of OPTN between the two centrioles was measured at the G1/S boundary and at a time point (4 h post-release) in S-phase. A clear preferential distribution of OPTN was observed at the G1/S boundary, followed by a more balanced distribution between the two centrioles as cells progressed into S phase (Fig. 5c, 5d, 5e).

**Fig. 5.**
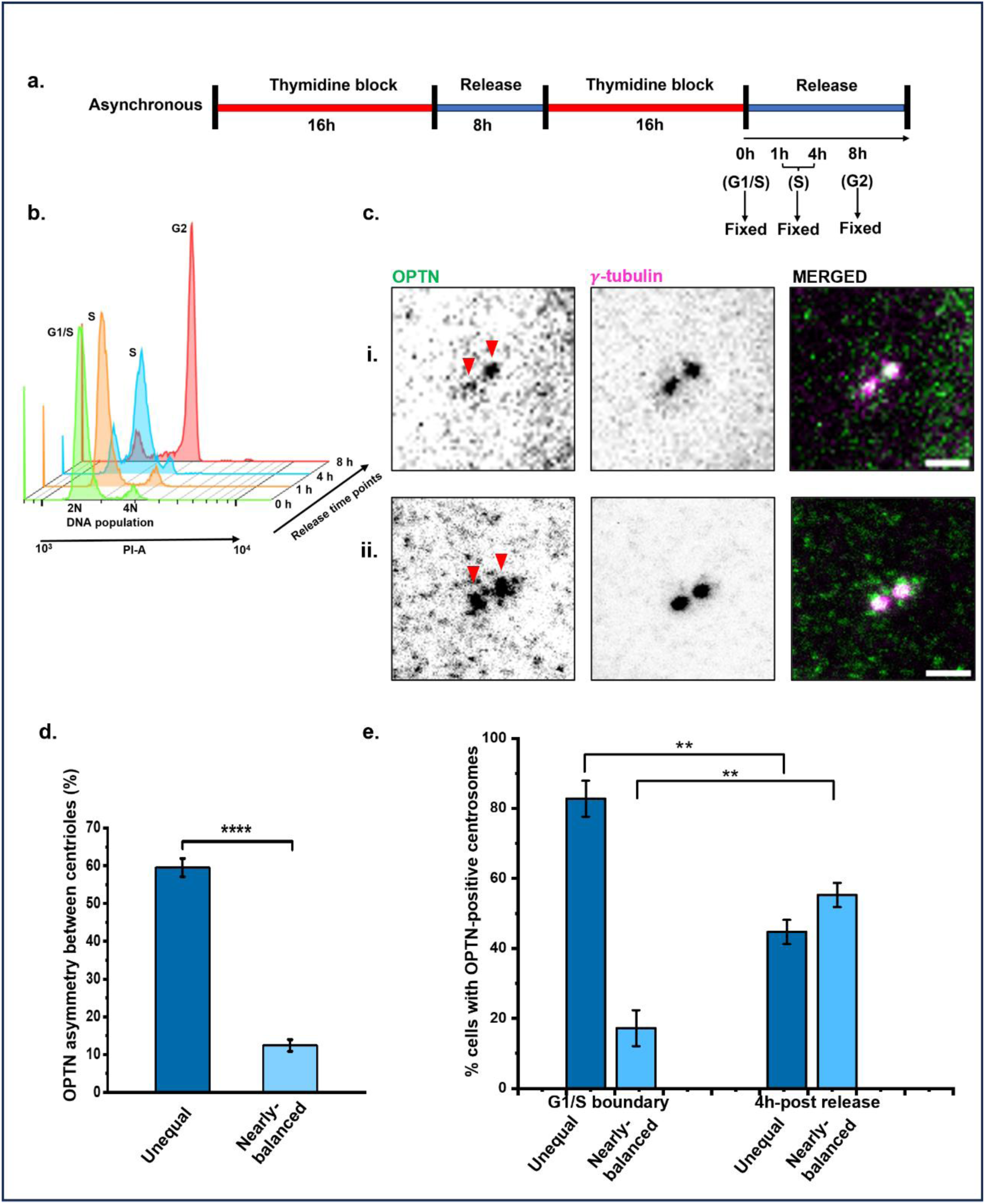
Asymmetric distribution of OPTN between centrioles during G1/S and S phases. (a) Schematic representation of the double-thymidine block following release to obtain an enriched population of cells at the G1/S boundary and in S phase. (b) Flow cytometry profiles of PI-stained HeLa cells synchronised as above. (c) Confocal images of cells co-stained for OPTN (green) and γ-tubulin (magenta) showing (i) an unequal and (ii) nearly balanced OPTN distribution between the two centrioles. Arrowheads indicate OPTN at the centrioles. Single channels are shown as inverted grayscale images. (d) OPTN asymmetry between centrioles was calculated as 100 × [(H − L)/H], where H and L represent the higher and lower CTCF values of centriolar OPTN, respectively. Data are presented as mean ± SEM (n = 30 cells per condition). (e) Quantification of the percentage of cells with unequal and nearly-balanced distribution of OPTN between two centrioles in G1/S and S phase (4 h post-release) cells, respectively. Data represent mean ± SEM from three independent experiments with 15-25 centrosomes analysed per experiment. Statistical analysis was performed using an unpaired two-tailed *t-test*, with significance defined as *p < 0.05, **p < 0.01, ***p < 0.001, ****p < 0.0001. Scale bars, 2 µm.

### 5. Centrosomal association of OPTN is maintained independently of microtubules

Having established the dynamic trafficking of OPTN in the centrosome-midbody axis, we next focused on its spatial organisation within the centrosome. To assess whether the centrosomal pool of OPTN requires an intact microtubule network for its retention, we first subjected cells to cold-induced microtubule depolymerisation. As per previous report of microtubule network depolymerisation through short cold shock, we incubated the HeLa cells on ice for 30 minutes ^33,34^. Cold induction, as an initial drug-independent approach, froze the cell’s dynamics and disrupted microtubules. Under this condition, we examined OPTN retention at the centrosome with reduced cellular dynamics. This allowed us to assess the pre-existing centrosomal OPTN pool following microtubule depolymerisation, minimising contributions from active recruitment or exit during the treatment.

Confocal microscopy of these cells revealed a disorganised mesh of tubulin structures, distinct from the well-defined filamentous microtubule network in control cells, confirming microtubular loss. Upon rewarming, rapid microtubule repolymerization was observed, confirming reversibility of the process (Fig. 6a). Interestingly, despite microtubule depolymerisation, co-staining of OPTN with γ- or β-tubulin separately, as well as the ratio of total OPTN intensity (CTCF) to γ-tubulin, showed no significant change in centrosomal OPTN intensity or its overall organisation relative to control cells (Fig. 6b, 6c, 6d, Supplementary Fig. S1). This suggests that centrosomal OPTN is stably maintained independent of microtubule integrity, possibly being anchored to other pericentriolar matrix scaffold proteins.

**Fig. 6.**
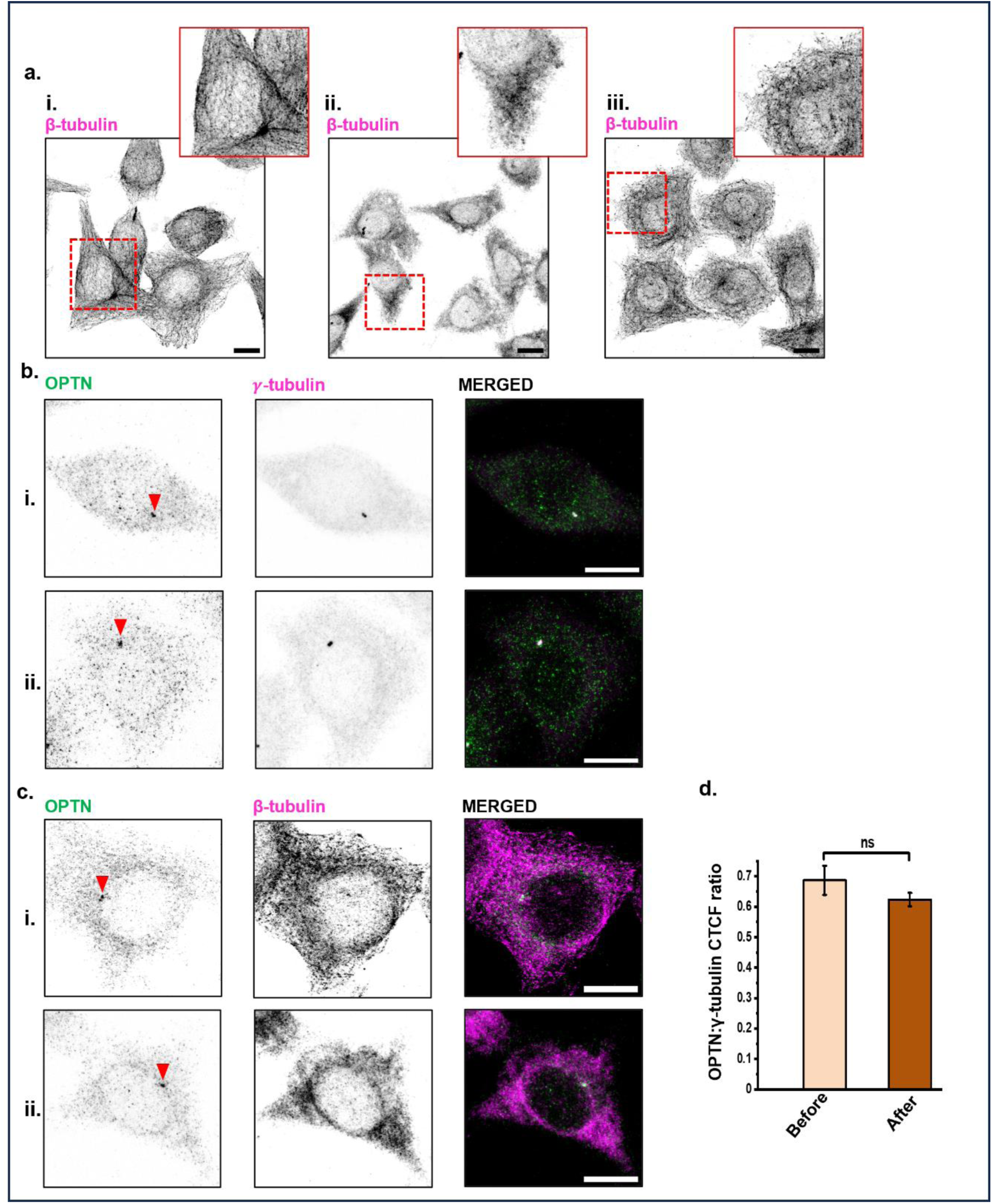
Retention of centrosomal OPTN following cold-induced microtubule depolymerisation. (a) Confocal images of microtubules probed with β-tubulin antibody in control cells (i), cold-treated cells (30 min) (ii), and heat-recovered cells (30 min) (iii) showing reversible microtubule depolymerisation and repolymerisation with temperature. Zoomed views (insets) are shown in the upper-right corner of each image. Confocal images of co-stained OPTN with (b) γ-tubulin and (c) β-tubulin, showing persistent localisation of OPTN at the centrosome before (b.i. and c.i.) and after (b.ii and c.ii.) microtubule depolymerisation. Arrowheads indicate OPTN at the centrosomes. Single channels are shown as inverted grayscale images. (d) Ratio of centrosomal total OPTN intensity to γ-tubulin (OPTN_CTCF_: γ-tubulin_CTCF_) was quantified for cells before and after microtubule depolymerisation. The data represent the mean ± SEM across n=30 cells for each condition. We performed statistical analysis using an unpaired two-tailed t-test, with significance defined as *p < 0.05, **p < 0.01, ***p < 0.001, ****p < 0.0001, and non-significance (ns) as p ≥ 0.05. Scale bars, 10 µm.

### 6. Dual regulation of centrosomal OPTN by microtubule-independent recruitment and microtubule-dependent exit

While retention of the OPTN pool at the centrosome under cold-induced static conditions strongly downplays any role for microtubules, the possibility of its active role in the recruitment dynamics of OPTN at the centrosome cannot be ruled out. To investigate, nocodazole, the microtubule-depolymerising agent, was used to disrupt microtubule integrity in HeLa cells and assess whether OPTN recruitment at the centrosome depends on microtubules. In this case, cells were treated with 0.5 μM nocodazole for 2 hours, then released into fresh medium. When observed with β-tubulin staining, depolymerised microtubules were visible after 2 hours of treatment, with a rapid recovery and polymerisation within 5 minutes of release (Fig. 7a). Besides, after 2 hours of treatment, OPTN exhibited an enlarged pericentriolar accumulation surrounding an intact γ-tubulin-defined centrosome (Fig. 7b, iii). This phenotype was distinct from the centrosomal signal of OPTN observed in the native/unperturbed and dimethyl sulfoxide (DMSO)-treated controls (Fig. 7b, i, ii). Quantification revealed a significant increase in the area and total centrosomal intensity of OPTN, as well as in the percentage of cells with an enlarged OPTN-positive pericentriolar pool following nocodazole treatment (Fig. 7c, 7d, 7e, 7f). The substantial accumulation of OPTN under a microtubule-depolymerised state strongly suggests a microtubule-independent recruitment of OPTN to the centrosome. Interestingly, when released (5 min), a rapid reversal of OPTN to its native-like signals was observed (Fig. 7b, iv). The imaging results were confirmed by quantifying the percentage of cells with centrosomal OPTN that returned to pre-treatment levels. (Fig. 7e, 7f).

**Fig. 7.**
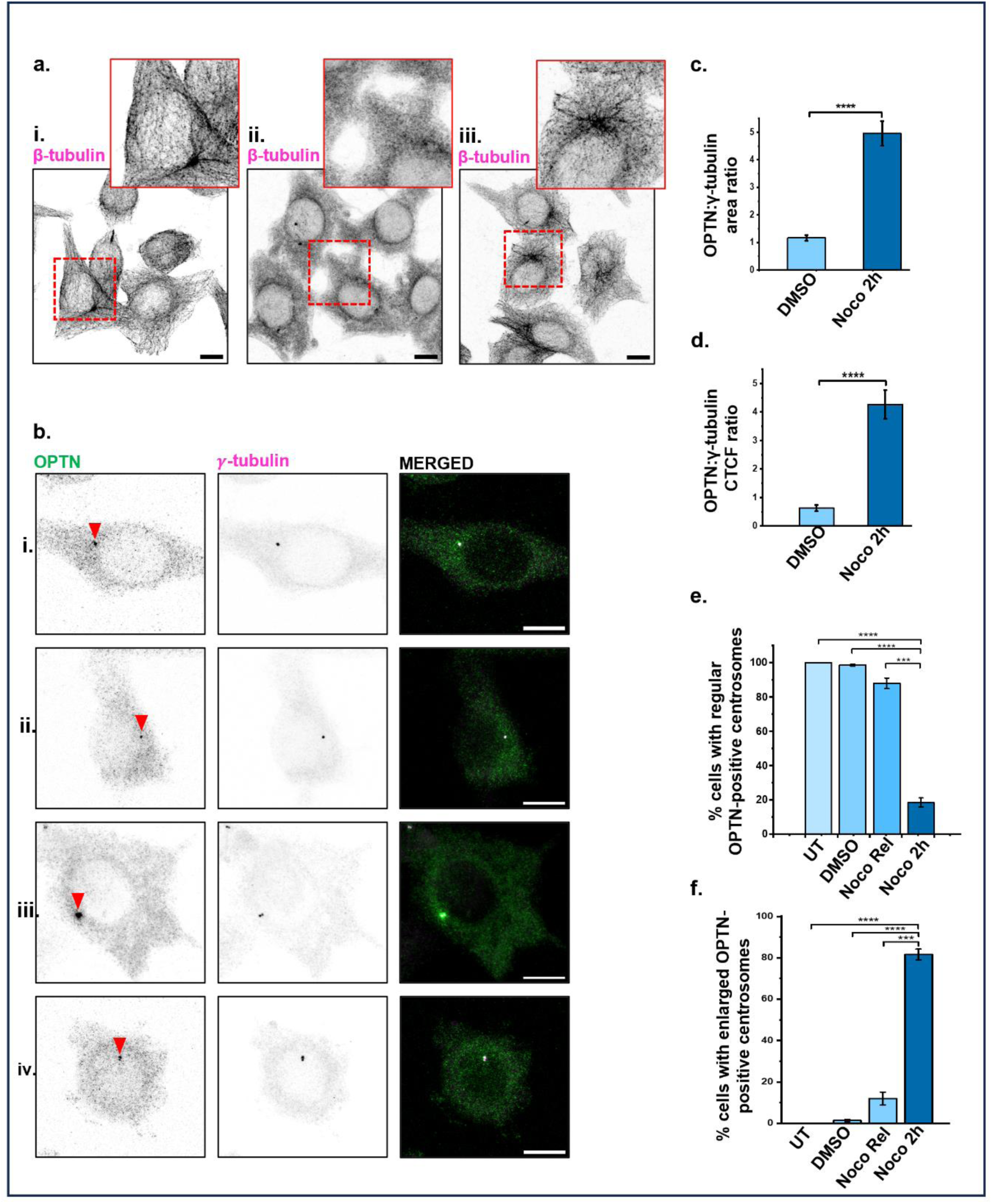
Microtubule-independent recruitment and microtubule-dependent exit regulate centrosomal OPTN levels. (a) Confocal images of cells stained for β-tubulin showing microtubule organisation under different conditions. (i) Filamentous, polymerised microtubules in control/unperturbed, native cells; (ii) Mesh-like, fragmented microtubules after 2-hour treatment with nocodazole (0.5 µM); (iii) Microtubule recovery within 5 minutes after release from nocodazole treatment. Zoomed views (insets) are shown in the upper-right corner of each image. (b) Images of cells co-stained with OPTN (green) and γ-tubulin (magenta). i) Untreated (UT) cell displaying a native centrosomal signal of OPTN. (ii) DMSO-treated (2h) cell as a control showing native-like centrosomal OPTN signal. (iii) Nocodazole-treated (2h) cell showing enlarged OPTN foci surrounding a γ-tubulin-marked centrosome. (iv) Recovered cell (5 min post-nocodazole washout) showing OPTN signal reversal to pre-treatment levels. Arrowheads indicate OPTN at the centrosomes. Single channels are shown as inverted grayscale images. The ratio of centrosomal OPTN area to γ-tubulin (OPTN_area_: γ-tubulin_area_) (c), as well as the ratio of centrosomal total OPTN intensity to γ-tubulin (OPTN_CTCF_: γ-tubulin_CTCF_) (d), was quantified for cells treated with 0.5 uM nocodazole and DMSO for 2 h. Data are presented for both (c) and (d) as mean ± SEM (n = 30 cells per condition). The percentage of cells with regular (e) vs enlarged pericentrosomal OPTN foci (f) was quantified under different conditions. Data are expressed as mean ± SEM for both (e) and (f) from three independent experiments, analysing 35-50 centrosomes per experiment. Statistical analysis was performed using an unpaired two-tailed *t-test*, with significance defined as *p < 0.05, **p < 0.01, ***p < 0.001, ****p < 0.0001. Scale bars, 10 µm.

Together, these observations indicate that transient accumulation of OPTN at the centrosome is a downstream effect of continuous microtubule-independent recruitment coupled with impaired microtubule-dependent exit of OPTN from the centrosome.

### 7. Radial dispersal of OPTN away from the centrosome on the microtubular track

Having evidenced OPTN accumulation at the centrosome under low-dose nocodazole, we investigated whether high-dose nocodazole treatment would give a different outcome. Therefore, a high concentration of nocodazole (33 µM) was used to examine whether the effect of microtubule depolymerisation on centrosomal OPTN dynamics depended on treatment dose and duration. Confocal imaging showed reduced microtubules after nocodazole-induced depolymerisation compared with control cells, with recovery upon release (Fig. 8a). HeLa cells were treated with 33 µM nocodazole for 30 minutes. OPTN accumulated at the PCM, resembling the low-dose, prolonged-treatment pattern, but the accumulation was brief and occurred within a limited time window (Fig. 8b, ii, Supplementary Fig. S2a). However, when analysed after 2 hours of treatment, most of the cells did not exhibit the prominent pericentrosomal accumulation of OPTN (Fig. 8b, iii, Supplementary Fig. S2b); instead, they remained largely confined to the centrosomal area, like the controls (Fig. 8b, i, vi).

**Fig. 8.**
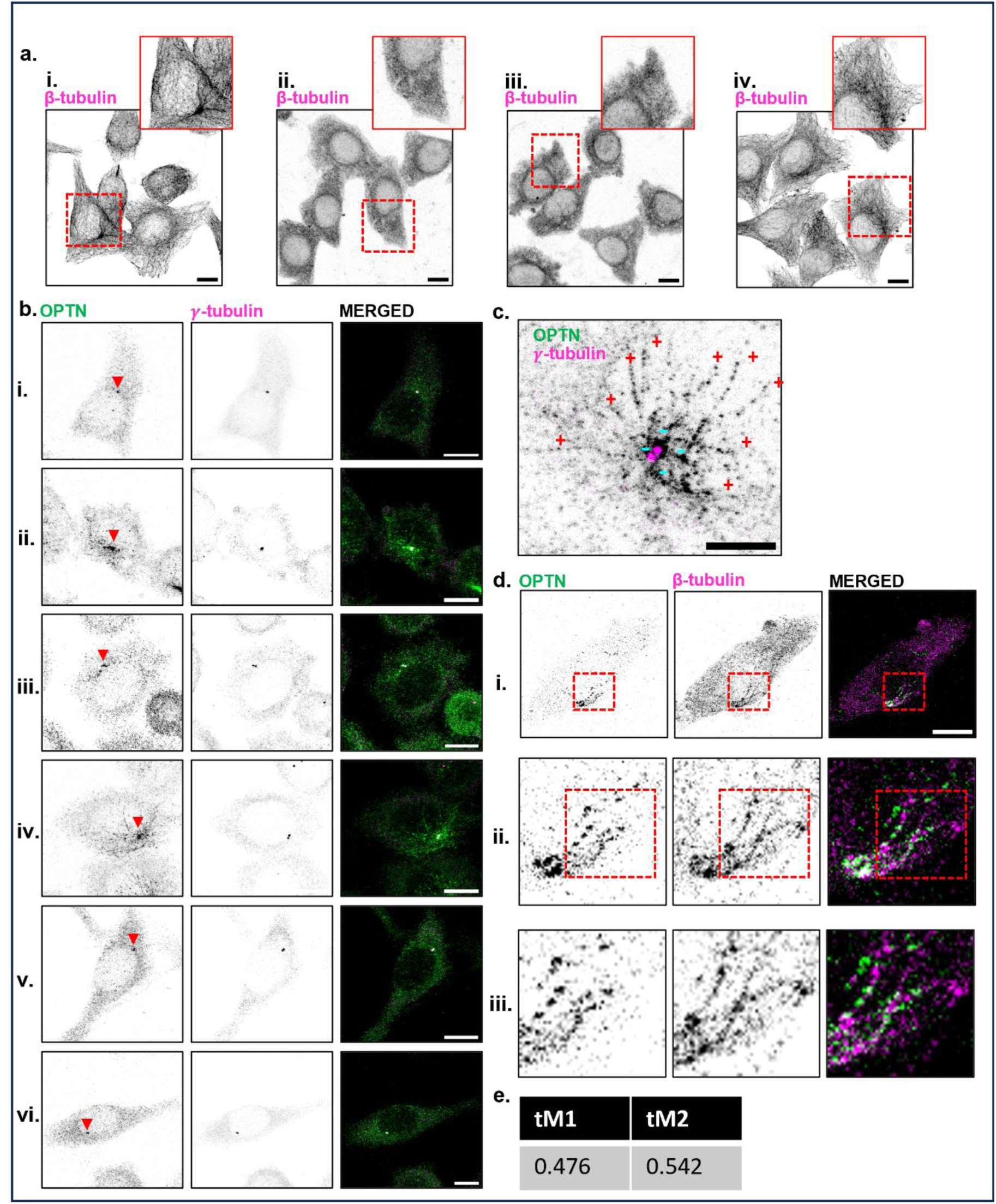
High-dose nocodazole-induced microtubule depolymerisation reveals time-dependent redistribution of OPTN. Immunofluorescence analysis of OPTN localisation in response to microtubule depolymerisation and regrowth. (a) Confocal images of microtubules probed with β-tubulin antibody in control cells (i), nocodazole-treated cells for 30 minutes (ii) and 2 hours (iii), and cells following washout (iv), confirming reversible microtubule depolymerisation and repolymerisation. (b) (i) In UT cells, OPTN (green) displays a compact pericentrosomal distribution surrounding the γ-tubulin-marked (magenta) centrosome. (ii) Short-term microtubule depolymerisation (33 µM nocodazole, 30 min) induces a pronounced accumulation of OPTN in an expanded, cloud-like structure around the centrosome, while γ-tubulin remains relatively compact. (iii) Prolonged nocodazole treatment (33 µM, 2 h) leads to a confined pericentrosomal pool of OPTN. (iv) Following the nocodazole washout after the 2 h treatment, cells were allowed to recover for 15 min. OPTN rapidly reaccumulates at the centrosome, showing a transient radial, star-like pattern emanating from the γ-tubulin-positive MTOC. (v) After 30 minutes of washing, OPTN displays the same pericentrosomal distribution as in the control. (vi) DMSO-treated cells for 2 h show a regularly confined OPTN signal at the centrosome, similar to UT cells. Arrowheads indicate OPTN at the centrosomes. Scale bar, 10 µm. (c) Radial appearance of OPTN (green) from the γ-tubulin-positive (magenta) MTOC towards the plus end of the extending microtubule. Scale bar, 5 µm. (d) Outward dissipation of OPTN (green) is partially tethered to β-tubulin-labelled (magenta) microtubules (i). The second (ii) and third (iii) rows display progressively zoomed-in images (red box) of the first row. Single channels are shown as inverted grayscale images. Scale bar, 10 µm. (e) Intensity data taken from the section (red box) of the second row showing 54.2 % of OPTN signal co-localised with β-tubulin. tM1 and tM2 are threshold Manders’ coefficients for channel 1 (β-tubulin) and channel 2 (OPTN).

Although prolonged treatment did not cause significant accumulation, a striking, radial, star-like distribution of OPTN outward from the γ-tubulin-marked centrosome was observed after 12-15 minutes of nocodazole washout (Fig. 8b, iv, 8c, 8d, Supplementary Fig. S2c). Colocalization of β-tubulin with OPTN showed partial alignment of OPTN’s radial structures with newly formed microtubules (Fig. 8d, 8e). However, at 30 minutes after recovery, when the microtubule network was fully reformed, OPTN returned to its usual centrosomal position (Fig. 8b, v).

## Discussion

In this study, we traced the association and dispersal path of OPTN between the centrosome and other macromolecular ensembles during cell cycle progression. Cell synchronisation and modulation of microtubule assembly/disassembly, combined with intensive confocal imaging, enabled systematic analysis of each stage. We report a cyclic centrosome-to-midbody repositioning of OPTN, followed by its subsequent retrieval to the centrosomes in the next cell cycle. Together with our previous identification of OPTN as a PCM-associated protein, these findings establish its dynamic centrosomal association, spatial organisation, and redistribution during cell cycle progression.

A central finding of this work is that OPTN is gradually redistributed from centrosomes to the midbody in a cell cycle-regulated manner during mitosis. Quantitative analysis of synchronised HeLa cells shows a progressive reduction in centrosomal OPTN from anaphase I to anaphase II, and it mostly disappears during late telophase to cytokinesis (Fig. 1). Importantly, this loss is observed only at the late stages of cell division, not during interphase, and is suggestive of an active dissociation of OPTN from the centrosome. We also observed a similar centrosome-to-midbody trafficking pattern in noncancerous HEK293 cells (Fig. 2), supporting the conserved nature of OPTN trafficking during cell division.

Building on these observations, we next discuss the detailed spatiotemporal dynamics and centrosomal regulation of OPTN in HeLa cells during cell-cycle progression. Following its release from centrosomes, OPTN undergoes a spatially ordered redistribution to cytokinetic structures. In this study, OPTN is first detected at the spindle midzone during late anaphase and early telophase, and over the course of cytokinesis, it progressively localises to the midbody. At the midzone, OPTN adopts a compact, disc-like organisation that is spatially distinct from the surrounding actomyosin contractile ring marked by NMIIB (Fig. 3). While Myosin-II effectively marks the contractile machinery in this study, future research involving actin co-staining will be very useful for better understanding the detailed structural interactions between OPTN and the broader actomyosin cortex.

Although the present work does not directly observe its dynamics along spindle microtubules, earlier analyses in asynchronous cell populations detected OPTN on the mitotic spindle ^27^. Taken together, these observations suggest a phenomenon in which spindle association represents a transient intermediate state, signifying directional anterograde transport of OPTN from centrosomes toward the midzone and midbody.

One plausible regulatory mechanism underlying this redistribution is post-translational modification (PTM). Mitotic PTMs are well defined as modulators of protein-protein interactions, subcellular localisation, and binding affinities during cell division [30-32]. PTM-dependent regulation could facilitate the release of OPTN from the PCM during mitosis and promote its association with midzone and midbody structures. Additionally, it can potentially do so via microtubule-based transport pathways [32]. Although the specific modifications involved remain to be identified, our results provide a logical framework for the tight stage-specific relocalisation of OPTN during mitotic progression. While our study establishes this foundational roadmap of OPTN trafficking, the precise upstream molecular mechanisms driving this cyclic orchestration remain to be fully elucidated. Future investigations utilising specific mitotic kinase inhibitors or phospho-deficient mutants will be essential to discover whether site-specific PTMs regulate this dynamic recruitment. Furthermore, identifying which specific PCM scaffold proteins directly anchor or recruit OPTN at the centrosome will provide critical mechanistic insights into how this spatial organisation is maintained independently of microtubules during recruitment, yet utilises them for anterograde transport.

Despite its disappearance from centrosomes during cytokinesis, OPTN is re-established gradually at the centrosomes following mitotic exit, suggesting that reorganised PCM scaffolds in G1 provide binding environments that are permissive for its centrosomal association (Fig. 4). This reversible association underscores the cyclical recruitment process of OPTN and indicates that its centrosomal presence is actively regulated during centrosome reassembly.

Moreover, the specific molecular anchors or interacting partners that mediate OPTN association with these macromolecular assemblies (centrosome or midbody) require thorough investigation. It is well established that OPTN directly interacts with key proteins such as Tank-binding kinase 1 (TBK1), Myosin VI, CYLD, and Polo-like kinase 1 (Plk1), all of which localise to the centrosome and have distinct regulatory roles during cell division and structural maintenance ^23,35–37^. TBK1 helps in spindle assembly, whereas Myosin VI, CYLD, and Plk1 support ciliogenesis, centriolar satellite proteostasis, and centrosomal maturation, respectively ^38–41^. It is highly plausible that the cyclic redistribution of OPTN mapped in this study is mediated by stage-specific interactions with these localised partners. Characterising which of these specific binding partners anchors the independent pool of OPTN at the PCM or mediates its handover to microtubule-based transport pathways is an essential next step in uncovering the full mechanistic landscape of OPTN during the cell cycle.

In our previous analysis of asynchronous cell populations, asymmetric distribution of OPTN between the two centrioles was described based on morphological and spatial criteria, such as early duplication and late S-phase stages ^27^. However, the lack of precise cell-cycle staging in asynchronous cells limited the temporal resolution. By synchronising cells in the present study, we refined this observation and demonstrated that preferential enrichment of OPTN at the mother centriole is established primarily during the G1/S boundary, coincident with daughter centriole separation (Fig. 5). This aligns with evidence suggesting that temporally regulated centriolar asymmetry is a defining feature of proteins associated with centrosome maturation and functional specialisation. The subsequent incorporation of OPTN at the daughter centriole during S phase further supports a maturation-coupled balancing program, consistent with its dynamic PCM localisation (Fig. 5).

A crucial insight from this research is that microtubules differentially regulate the centrosomal recruitment, retention, and exit of OPTN. This became evident from the cold-induction and nocodazole-mediated microtubule disruption studies.

Assuming the association of OPTN at the centrosome is microtubule-dependent, it would have accompanied the cold-induced disruption. In contrast, these cells displayed an intact OPTN pool at the centrosome, possibly through association with PCM scaffolds, and were unaffected by microtubular instability (Fig. 6).

In the case of nocodazole treatment (using a low 0.5 μM concentration for 2 hours), which caused reversible depolymerisation of microtubules, a rapid accumulation of OPTN around centrosomes, aligned with the formation of an enlarged PCM, was observed (Fig. 7b, iii). Given that this recruitment occurs under microtubule-depolymerised conditions, it supports an alternative scaffold-mediated assembly at the centrosome. In a way, this recruitment process resembles the recruitment of proteins such as Aurora A Kinase, Plk1, and the type one protein phosphatase to the centrosome by a scaffolding protein, CEP192 ^14,42^.

Interestingly, upon immediate nocodazole washout, the accumulated OPTN signal disperses and returns to its basal centrosomal levels. This rapid recovery indicates that the structure is highly transient and dynamic, and, unlike recruitment, the exit of OPTN from the centrosome is strongly dependent on microtubule integrity (Fig. 7). Thus, under depolymerised conditions, OPTN appears to be continuously recruited but unable to efficiently exit from the centrosome, resulting in its transient pericentrosomal accumulation. Interestingly, centrosomal OPTN levels matched when cells were blocked either for a short time (∼30 minutes) with a high dose of nocodazole (33 µM) or for a longer time (2 hours) with a low dose of nocodazole (0.5 µM) (Fig. 7b, iii; Fig. 8b, ii). These concentrations allowed us to examine the OPTN response across different durations and levels of nocodazole exposure. However, extending the blocking time (2 hours) at higher concentrations led to OPTN levels like those of UT controls (Fig. 8b, iii). We reasoned that extended treatment with higher concentrations of nocodazole not only depolymerises microtubules but also induces acute stress. The stress could trigger alternative clearance processes (autodegradation, proteasomal degradation), allowing rapid recycling of accumulated OPTN away from the centrosome (Fig. 8b, iii). A particularly striking phenomenon was observed when the cells were washed free of high-dose nocodazole after 2 hours and released for 12-15 minutes. OPTN emerges radially from the centrosome, apparently latched onto nascent microtubule tracks (Fig. 8b, iv, 8c, 8d). Such association of OPTN with microtubules, along with other adapter proteins, has been previously shown for kinesin-mediated anterograde transport ^18^.

Images taken at a later time point (30 minutes post-nocodazole wash) showed the disappearance of OPTN-bound microtubular tracks and the restoration of centrosomal OPTN to normal levels (Fig. 8b, v). This indicates that OPTN undergoes a precisely time-regulated, transient spatio-temporal association with the microtubule during its exit from the centrosome.

Collectively, our findings support a model in which OPTN functions as a dynamic PCM-associated factor whose localisation is governed by cell cycle stage and centrosomal architecture-linked transport. This model further suggests that centrosomal OPTN levels are maintained by a dynamic balance between its scaffold-mediated recruitment at the centrosome and microtubule-dependent exit. Importantly, OPTN also exhibits clear cell-cycle-dependent dynamics. During mitosis, OPTN is progressively released from centrosomes and transferred toward cytokinetic structures, passing through transient intermediate states before incorporation into the midzone and midbody. After mitotic exit, OPTN re-associates with reassembled centrosomal structures in G1 of the subsequent cell cycle, initially showing preferential enrichment at the mother centriole at the G1/S boundary before achieving a balanced distribution as duplication proceeds. This study also shows that OPTN distribution is tightly linked to precise, narrow windows of the cell cycle, allowing it to serve as a marker for specific cell-cycle stages.

Ideally, a direct functional assignment linking the downstream consequences of optineurin (OPTN) disruption at these designated sites would have further illuminated the field. However, this study is an exploratory effort that provides detailed spatiotemporal mapping of endogenous OPTN, highlighting cyclic trafficking pathways, specific stages of division, and precise routes of macromolecular assembly. We believe that this revelation will greatly reduce the experimental search space by providing a focused platform for researchers to dissect the site-specific molecular functions of OPTN at the centrosome, spindle and midbody.

To our knowledge, this is the first systematic representation of OPTN trafficking dynamics across cell-division assemblies in the context of cell cycle progression. These findings may also provide a new perspective on the disease-associated roles of OPTN. Several OPTN mutations have previously been associated with NDs such as ALS and POAG, while emerging evidence also implicates aberrant cell division in neurodegeneration ^24,25,43^. Our report bridges these findings and opens avenues to explore disease-related mutants with aberrant OPTN trafficking. In other words, whether the clinical mutants of OPTN correlate with its aberrant dynamic distribution during the cell cycle could be an interesting study, providing unique insights into the pathogenic processes of these disorders.

## Materials and methods

### 1. Cell line maintenance and Reagents

HeLa and HEK293 cells were maintained in Dulbecco’s modified Eagle’s medium (DMEM, Gibco) supplemented with 10% (v/v) fetal bovine serum (FBS, Gibco) and 1% (v/v) penicillin-streptomycin (Gibco). Cells were cultured at 37 °C in a humidified incubator with 5% CO_2_. Trypsin (Gibco) was used to dissociate cells. DAPI (SRL) was used for nuclear staining. ProLong glass antifade mounting agent (PGAMA) was purchased from Invitrogen.

### 2. Antibodies

The primary antibodies used for the immunofluorescence studies were rabbit anti-optineurin (1:250, CST, E4P8C-monoclonal, raised against full-length human OPTN), mouse anti-γ-tubulin (1:500, Thermo Fisher Scientific; MA1-19421, recognizes C-terminal amino acids from 434 to 449 in human γ-tubulin, and the epitope was located in the amino acid sequence TRPDYI), mouse anti-β-tubulin (1:200, ABclonal, Cat# AC021, raised against recombinant human β-tubulin) and mouse anti-MYH10 (1:200, Santa Cruz, Cat# sc-33729, raised against amino acids 1964-1976 of purified myosin heavy chain of human origin to detect NMIIB). Alexa Fluor 488 goat anti-rabbit (1:1000, Invitrogen, Cat# A-11070) and Alexa Fluor 568 goat anti-mouse (1:1000, Invitrogen, Cat# A-11019) antibodies were used as secondary antibodies for immunofluorescence.

### 3. Cell synchronisation

Thymidine (Sigma) was used to synchronise HeLa and HEK 293 cells, and Nocodazole (MedChemExpress) was used to synchronise HeLa cells. Cells growing in 6-well or 24-well plates were treated with thymidine at 2.5 mM per well, a concentration widely reported to efficiently induce G1/S synchronisation, and incubated for 16 hours in a humidified incubator at 37 °C ^44^. Thymidine was removed after 16 h, and cells were washed thrice with 1X phosphate-buffered saline (PBS). Fresh, pre-warmed media were added, and the cells were incubated for an additional 8 h. A second thymidine treatment to a final concentration of 2.5 mM was given, and the cells were incubated for an additional 16 h at 37 °C. Cells at the G1/S boundary after the second thymidine treatment were washed 3 times with pre-warmed 1X PBS and incubated in pre-warmed fresh medium for release. Cells were collected at subsequent time points, followed by DNA staining with PI for cell cycle analysis Cells were synchronised at the G2/M boundary using nocodazole-mediated mitotic arrest. Briefly, cells at ∼70-80% confluency were treated with nocodazole (75 ng/mL), within the reported effective range of 50-100 ng/mL, for 18 h at 37 °C ^32,45^. Following treatment, mitotic cells were collected by mechanical shake-off, exploiting their rounded morphology and reduced adhesion. Collected cells were washed thrice with pre-warmed complete medium by centrifugation to remove residual nocodazole. The harvested mitotic population was subsequently reseeded in fresh nocodazole-free medium and allowed to progress synchronously through mitosis and into the next cell cycle for downstream experiments.

### 4. Microtubule depolymerisation by nocodazole treatment

Cells were grown on glass coverslips and treated with nocodazole dissolved in DMSO (HiMedia) for 30 mins or 2 hours at 37 °C. Nocodazole was then removed by briefly washing the cells in 1X PBS. Cells were then allowed to recover in growth medium at 37 °C. After recovery, cells were fixed with methanol at subsequent time points for 3 min at −20 °C. Cells exposed to DMSO (v/v) served as the control.

### 5. Microtubule depolymerisation by cold treatment

To induce microtubule depolymerisation, cells grown on coverslips were subjected to cold treatment by adding 4-(2-hydroxyethyl)-1-piperazineethanesulfonic acid (HEPES) buffer to the complete medium and incubating on ice for 30 minutes, as previously reported ^34^. Following cold exposure, cells were either immediately fixed to assess microtubule loss or released into warm complete medium (37 °C) for another 30 minutes to allow microtubule regrowth.

### 6. Flow Cytometry

Cells were trypsinised, washed twice with 1X PBS, fixed with 75% pre-chilled ethanol by dropwise addition under continuous vortexing, and incubated on ice for 30 minutes. Fixed cells were washed thrice with 1X PBS by centrifugation. The cell pellet was then resuspended in fresh 1X PBS containing 20 μg/ml RNase A and incubated for 30 minutes at room temperature. Next, a 10 μg/ml PI solution was added to the cell solution, and the mixture was incubated for 5 minutes at room temperature. Cellular DNA content was analysed using a BD Fortessa X20 Flow Cytometer with the 561 nm laser, and acquisitions were analysed using BD FACS DIVA 9. For each sample, at least 10,000 cells were counted in triplicate. Representative graphs were prepared using FlowJo v11.2 software.

### 7. Immunofluorescence

HeLa cells were grown on a glass coverslip and fixed with pre-chilled methanol for 3 min at - 20°C at the defined time points. After drying, coverslips were rehydrated in 1X PBS for 15 min at room temperature. Cells were incubated with primary antibodies overnight at 4 °C. Following three rinses with 1X PBS, the cells were incubated with Alexa Fluor-conjugated secondary antibodies (Alexa Fluor 488 or 568) for 1 h at room temperature. After rinsing three times with 1X PBS, cells were incubated with DAPI (1 μg/mL) for 5 min at room temperature, then washed three times with 1X PBS and mounted on glass slides using PGAMA for microscopy. All antibodies were diluted using 0.2% bovine serum albumin (BSA).

### 8. Imaging

Confocal images were acquired with a 63x/1.3 oil objective using a Leica TCS SP8 confocal laser scanning microscope (CRF, at IIT Delhi), equipped with 405 nm, 488 nm, and 552 nm lasers. Image acquisition was done using the LASX application software.

### 9. Image processing and quantitative image analysis

Images were processed using Fiji software. For quantitative image analysis, images were acquired with no pixel saturation to ensure accurate intensity measurements.

#### Quantitative image analysis

Corrected mean fluorescence = Mean Fluorescence - Mean Background Fluorescence This was calculated separately for OPTN and γ-tubulin

**Optineurin:γ-tubulin corrected mean ratio** = OPTN_corrected mean_: γ-tubulin_corrected mean_ CTCF = Integrated Density - (Area x Mean Background Fluorescence)

This was calculated separately for OPTN and γ-tubulin

**Optineurin:γ-tubulin CTCF ratio** = OPTN_CTCF_: γ-tubulin_CTCF_

**OPTN asymmetry between centrioles** = 100 x [(H-L)/ H]

where H and L represent the higher and lower centriolar OPTN CTCF values, respectively.

### 10. Statistical analysis

Statistical significance was computed using an unpaired two-tailed *t-test* for the various data sets in MS Excel. The graphs were plotted using MS Excel or OriginPro 2020b software. Data are presented as mean ± SEM. For analyses that were expressed as percentages of cells, three independent experiments were performed, with each experiment contributing a single value. For the remaining quantitative measurements, individual cells or centrosomes were treated as individual measurement units (n), with each cell or centrosome contributing a single value.

## Supplementary Information

Attached at the end.

## Acknowledgements

A.C. and K.K. acknowledge the fellowship support given by IIT Delhi. All the authors acknowledge the CRF IIT Delhi for the confocal imaging facility (Cell Biology Lab, Sonipat) and for flow cytometry. A.C. and K.K. thank Prof. Archana Chugh for providing the HeLa cell line, Prof. Tapan Kumar Nayak for providing the HEK 293 cell line, and Prof. Anita Roy for providing some of the antibodies. Prof. Chinmoy S. Dey is hereby acknowledged for his critical scientific input.

## Author Contributions

Ankan Chakraborty and Bishwajit Kundu contributed to the study conception and design. Material preparation, data collection and analysis were performed by Ankan Chakraborty and Koyena Kundu. The first draft of the manuscript was written by Ankan Chakraborty, Koyena Kundu, Bishwajit Kundu, and Dominique Weil, and all authors commented on previous versions of the manuscript. All authors read and approved the final manuscript.

## Funder Information Declared

This work has not been funded by any specific funding agency.

## Data availability

All the data will be made available from the corresponding author upon reasonable request.

## Declaration of generative AI and AI-assisted technologies in the writing process

AI-assisted technologies [ChatGPT (OpenAI) and Grammarly] were used sparsely to enhance the language and readability of the manuscript. After employing these tools, the authors reviewed and edited the content as needed and took full responsibility for the publication’s content.

