## Supplementary figures and images for "Tracing the dynamic recruitment trajectory of Optineurin between the centrosome and other macromolecular assemblies, signifying its role in cell cycle progression"

### Supplementary Fig. S1

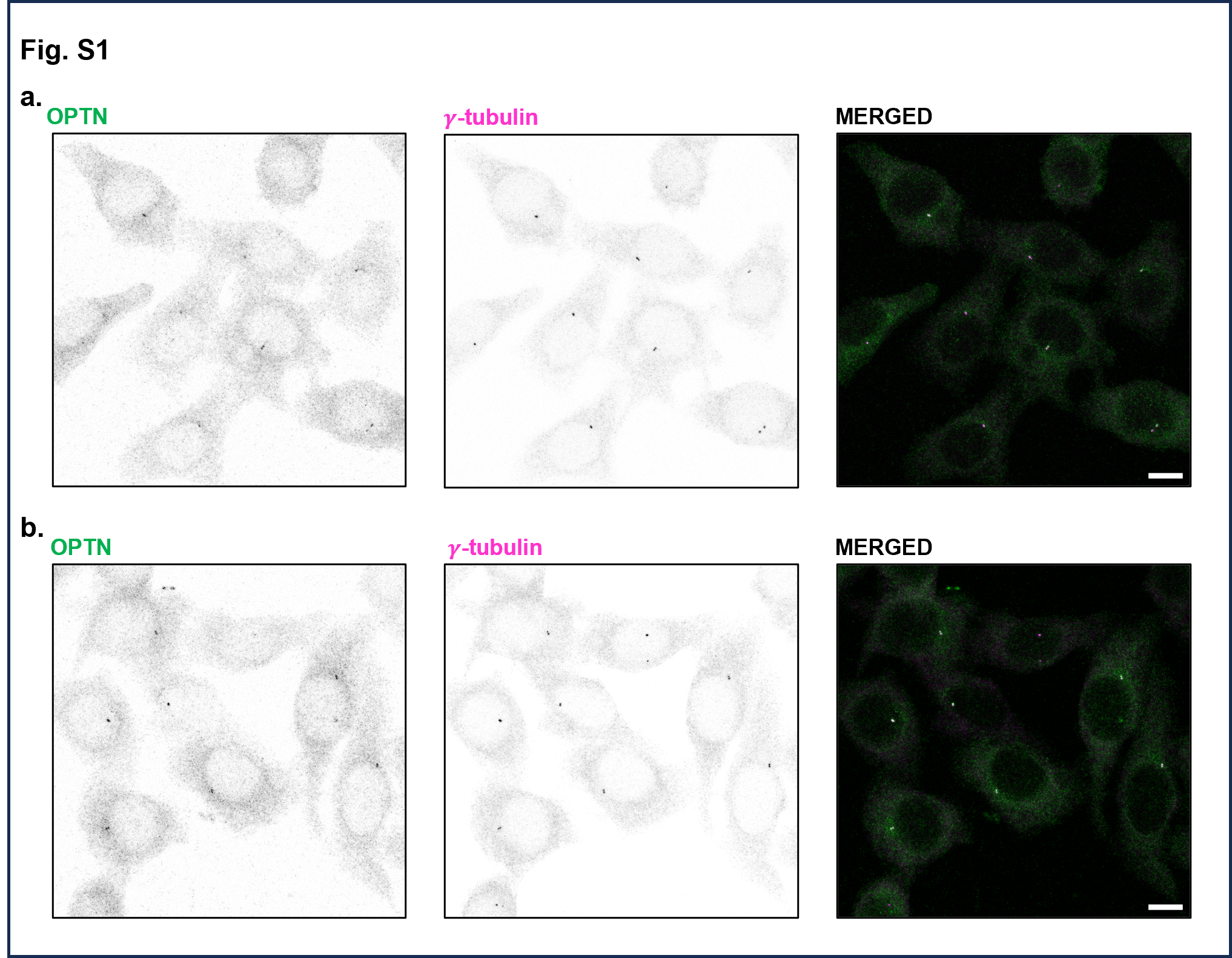

### Supplementary Fig. S2

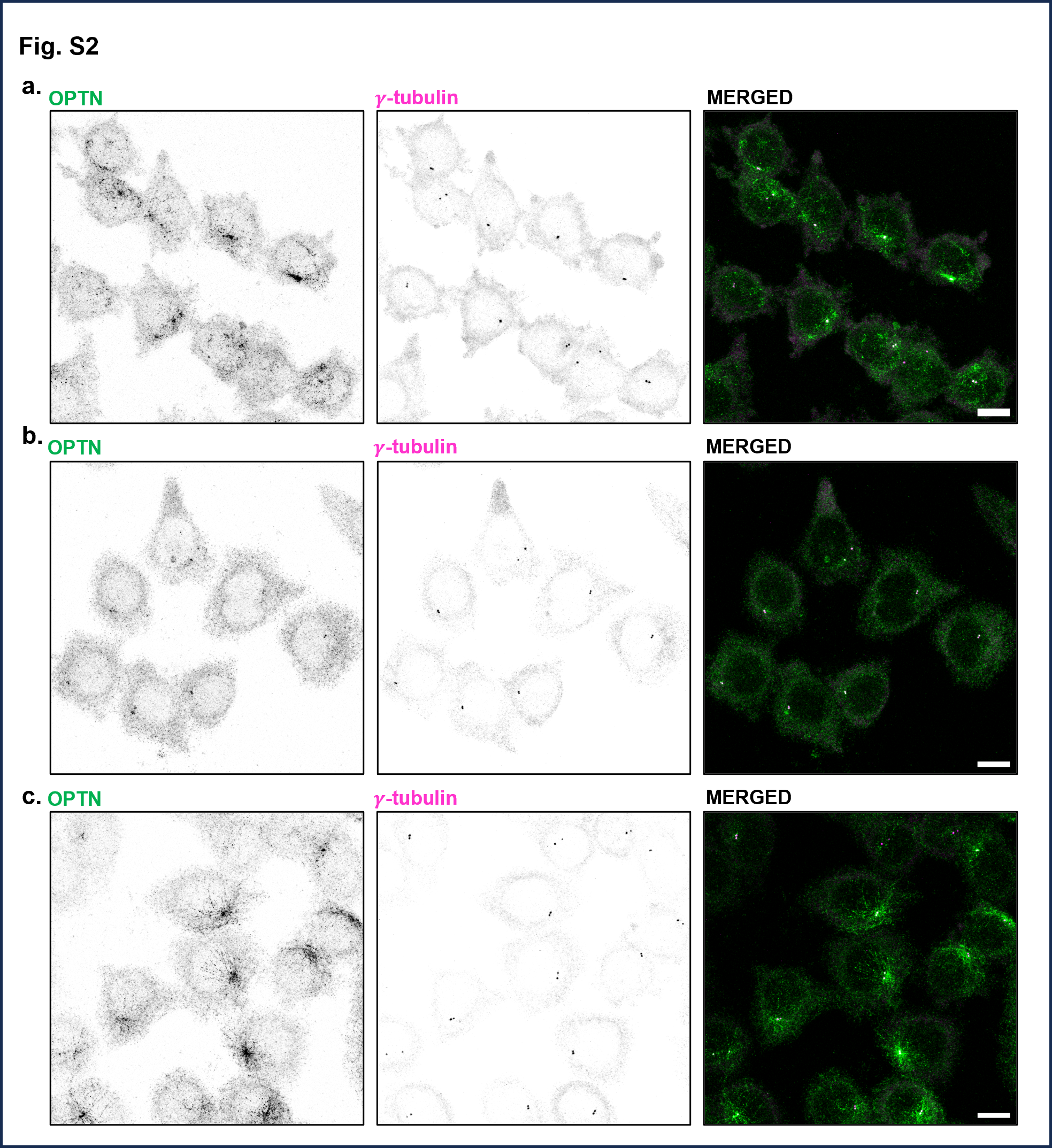
