## Supplementary Figure legends for "Tracing the dynamic recruitment trajectory of Optineurin between the centrosome and other macromolecular assemblies, signifying its role in cell cycle progression"

**Fig. S1. OPTN is retained at the centrosome following cold-induced microtubule depolymerisation.** Field image of cells co-stained for OPTN (green) and γ-tubulin (magenta) displaying continuous presence of OPTN at the centrosome before (a) and after (b) microtubule depolymerisation. Scale bars, 10 µm.

**Fig. S2.** **OPTN redistribution pattern following a high-dose nocodazole-induced microtubule depolymerisation**. Field image of short-term (33 µM for 30 min) nocodazole-treated cells showing (a) an accumulation of OPTN (green) in an expanded structure around the centrosome, (b) cells treated with nocodazole for long-term (33 µM for 2 h) showing a regular pericentrosomal OPTN pool, (c) cells washed out after the 2 h nocodazole treatment (33 µM for 2 h) and allowed to recover for 15 min, showing radial, star-like OPTN dissipating out from the γ-tubulin-positive MTOC (red). Scale bars, 10 µm.
